# Comparative genomics of Japanese encephalitis virus shows low rates of recombination and a suite of sites under episodic diversifying selection

**DOI:** 10.1101/2023.06.15.545193

**Authors:** Mark Sistrom, Hannah Andrews, Danielle Edwards

## Abstract

Japanese encephalitis virus (JEV) is the dominant cause of viral encephalitis in the Asian region with 100,000 cases and 25,000 deaths reported annually. The genome is comprised of a single polyprotein that encodes three structural and seven non-structural proteins. We collated a dataset of 347 complete genomes from a number of public databases, and analysed the data for recombination, evolutionary selection and phylogenetic structure. There are low rates of recombination in JEV, subsequently recombination is not a major evolutionary force shaping JEV. We found a strong overall signal of purifying selection in the genome, which is the main force affecting the evolutionary dynamics in JEV. There are also a small number of genomic sites under episodic diversifying selection, especially in the envelope protein and non-structural proteins 3 and 5. Overall, these results support previous analyses of JEV evolutionary genomics and provide additional insight into the evolutionary processes shaping the distribution and adaptation of this important pathogenic arbovirus.

**Author Summary:** This comparative study of Japanese Encephalitis Virus is the largest genomic analysis of the virus to date. We undertake a suite of analyses to investigate phylogenetic relationships, rates of recombination and patterns of genomic selection. We show that recombination is not a significant driver of evolution in JEV, demonstrate support for previous phylogenetic reconstructions of the virus, and find a number of sites across the genome under episodic diversifying selection. These adaptive hotspots of evolution serve as key genomic points for the adaptive evolution of this important vector borne pathogen.

## Introduction

Japanese encephalitis virus (JEV) is an arbovirus belonging to the Flaviviridiae family with a zoonotic cycle involving swine as reservoir hosts, waterbirds as carriers and mosquitoes of the two genera *Culex* and *Aedes* as vectors (1, 2). While humans are dead-end hosts for JEV as they generally display low viremias insufficient to allow for infection of feeding mosquitoes (3), JEV infections of humans have significant health implications, with around 100,000 cases of human JEV annually (4, 5) resulting in approximately 25,000 deaths (6). While only between 0.1-4% of human infections result in symptoms (7), symptomatic cases have a fatality rate of 20-30% (8) and 30-50% of survivors develop long term neurological/psychiatric sequelae (9). Despite the existence and use of several safe and effective JEV vaccines (8), JEV remains the dominant cause of viral encephalitis in the Asian region – meaning that understanding the evolutionary driving forces governing the range and pathogenicity of JEV are critical to ongoing management and control of this neglected tropical disease.

The JEV genome is 11kb, positive sense, single stranded RNA that comprises a single open reading frame encoding a large poly protein that is co- and post-translationally cleaved into three structural proteins – capsid (C), precursor to membrane (prM) and envelope proteins (E) and seven non-structural, accessory proteins (NS1, NS2A, NS2B, NS3, NS4A, NS4B, NS5) (10, 11). The NS1 protein plays an essential role in genome replication (12) – mutations within this gene can have marked effects on RNA replication and infectious virus production (13). NS2a and b are small, membrane associated proteins that play a role in virus assembly, RNA replication and interferon inhibition (14, 15). Mutations in NS2a have been shown to block virus assembly (16). NS3 is a large, multifunction protein, encoding enzymatic activities necessary for polyprotein processing, RNA replication, virus assembly and apoptosis (16–18). NS4a and b are both small, hydrophobic proteins. NS4a is involved in RNA replication via a genetic interaction with NS1 (19), and can induce membrane rearrangements and/or the formation of autophagosomes (20). Mutations in NS4a confer resistance to flavivirus RNA replication inhibitors (21). NS4b co-localizes with NS3 at sites of RNA replication, and is involved in blocking interferon signalling (22). NS5 is large multifunctional protein involved in RNA capping and RdRP activities, as well as the induction of interleukin-8 secretion and blocking interferon signalling (23, 24).

There are 5 recognized evolutionary lineages of JEV (GI-V) (25). Historically, GIII was the dominantly detected strain, however it has recently been superseded by GI (25–27). GII is largely confined to Southeast Asia and Northern Australia (28), and GIV and GV are generally confined to tropical Southeast Asia (29), however the 2022 outbreak of JEV in Northern Australia was determined to be GIV, representing a range expansion of this genotype (30). Vaccines are largely derived from GIII genotypes (29, 31) and a growing body of evidence suggests that these vaccines show reduced efficacy toward GI and GV strains (32–34).

JEV is an evolutionarily dynamic pathogen with fluid transmission parameters associated with variations in host/vector range and climate change (35, 36). Further, virulence, pathogenicity and immunogenicity appear to vary between strains and remain in flux as the virus evolves in the face of exceptionally dynamic environmental factors (26, 35). Resultantly, contemporary comparative genomic studies are necessary to better understand and predict epidemiological patterns of JEV.

In this study, we analyse the complete genomes of 347 JEV isolates, identifying patterns of polymorphism, phylogeny, recombination and selection. We find that while the genome is predominately under purifying selection, there are several siteswhich are subject to adaptive evolution across the phylogeny of JEV. It is likely that the adaptive evolutionary processes underlying the observed dynamism in the host, vector and geographic range of JEV, along with changes in virulence, pathogenicity and immunogenicity are being driven by a relatively small number of mutational changes at the genome scale.

### Materials and Methods

We downloaded seven isolate genomes from the NCBI Short Read Archive (SRA) using SRAtoolkit v3.0.0 (37) and one assembly from the NBCI Assembly database. A further 356 complete genomes were downloaded from the NCBI Nucleotide database using Batch Entrez (38). A further 36 samples were retrieved from the DNA Databank of Japan (DDBJ) SRA database (39), and a further 356 complete genomes were downloaded from the DDBJ nucleotide database (39), for a total dataset of 756 records. When filtered for redundancy, record accuracy and length, a sequence set of 347 sequences were selected for further analysis. SRA genomes were filtered for read quality using Trimmomatic v0.32 (40) under default parameters, and aligned to a serotype O reference strain (GCA_000863325.1) using BWA v0.7.17 (41) with the mem function. Output SAM files were then sorted and converted to BAM format using SAMtools v1.17 (42). Variant detection of each BAM file was undertaken using the mpileup function of BCFtools v1.17 (42) before being exported in FASTA format using the consensus function. SRA, Assembly and nucleotide data were then aligned in FASTA format using MUSCLE v5 (43) using the super5 algorithm and guide tree permutation enabled.

Recombination across the genome was calculated using the program RDPv5 (44) with default settings, which implements several methods to detect recombination in a given sequence alignment. Phylogeographic reconstructions were undertaken for the whole genome as recombination was not found to have a significant impact on genome structure using RaxML (45). Trees were estimated using a GTR model with 1000 bootstrap replicates. Selection was initially evaluated for the whole genome by calculating nucleotide diversity (π) and (θ) using a sliding window analysis with a window length of 100 and step size of 25 sites implemented in DNAsP v6 (46). Selection for each gene was tested initially using a codon-based Z test of neutrality implemented in MEGA11 (45). We calculated overall average Dn-Ds for each gene and probability of neutral model fit using the Nei-Gojobori method with 500 bootstrap replicates. Missing sites were treated with partial deletion with a cut-off of 95%. We further evaluated selection using a number of analyses implemented in the HyPhy2.5 (48). We implemented a Branch-site Unrestricted Statistical Test for Episodic Selection (BUSTED) (49) to evaluate each gene for episodic selection using the tree calculated earlier along all branches of the phylogeny. Secondly, a Mixed Effects Model of Evolution (MEME) (50) was used to test each gene for specific sites under diversifying selection using the same tree as a guide.

## Results and Discussion

### Recombination

The results of recombination analyses are reported in Table 1 and Figure 2. We found limited evidence of recombination in our dataset, especially in comparison with other viruses for which recombination is noted to be a driving evolutionary force (51–53). Recombination is not widely reported in unsegmented arboviruses (54), however recombination via template switching is thought to be an important driver of evolution in RNA viruses (55) and thought to be most likely to occur within vertebrate hosts during multi-strain infections (56). Recombination of JEV would have significant potential implications for vaccine efficacy. We did find evidence of a small number (n =18) of recombinant strains from both GI and GIII isolates, largely between closely phylogenetically related isolates and in the mid region of the E protein and NS5 protein respectively (Table 1, Figure 2a). One instance of recombination between a GI and GIII strain was detected, but the support for this recombination event was only by two of five methods implemented by RDP5 (44) and comprised of a short fragment in the 3’ end of the genome, which contains the most missing data in our alignment. Resultantly, this result is low confidence. We therefore conclude that recombination during multi-strain infection is unlikely to be a major driver in the evolution of genomic diversity in JEV.

**Table 1:**
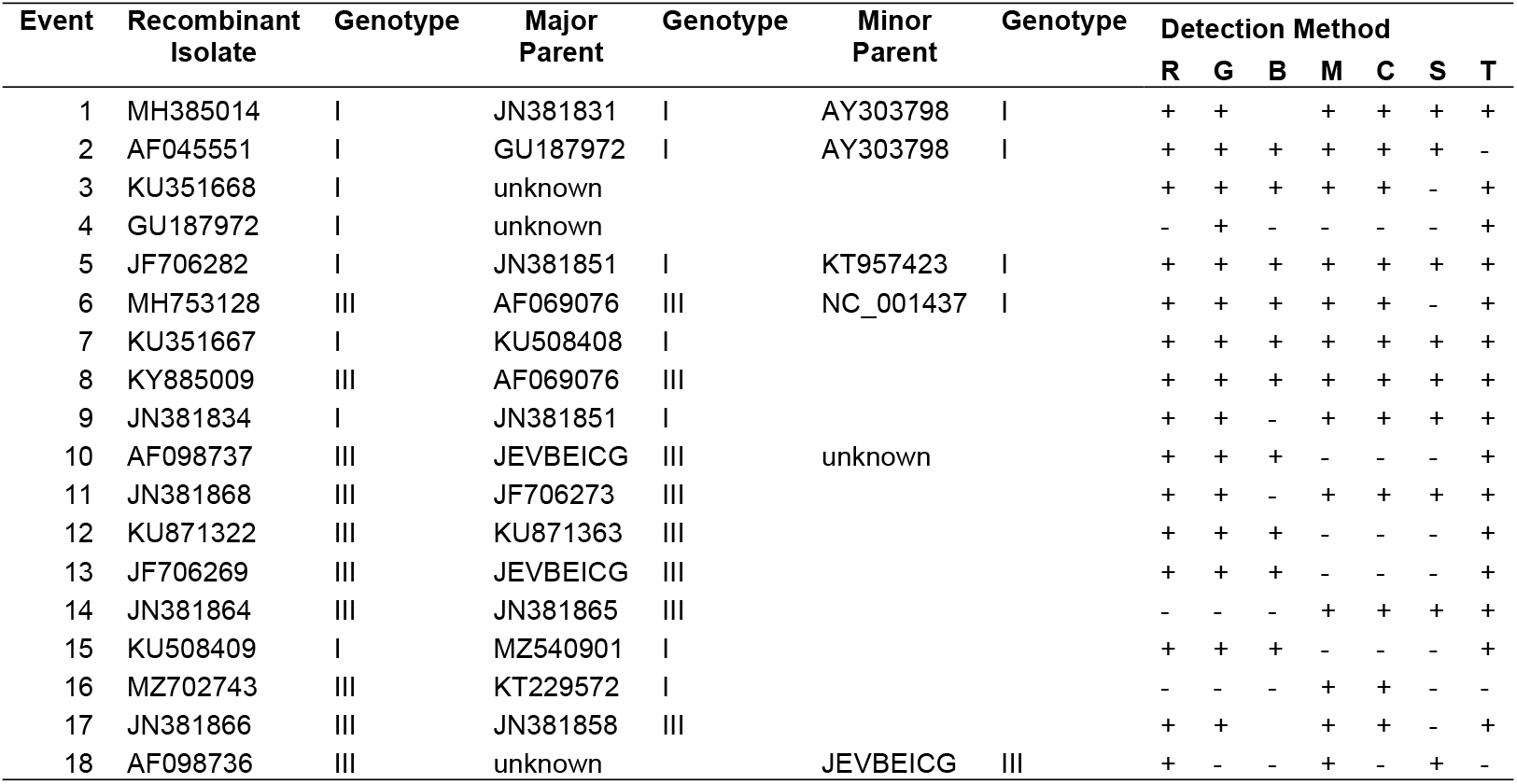
Recombinant strains as detected by RDP5 analysis. All events were between isolates of the same genotype with the exception of those with unknown parents, and event 16. R – RDP, G – GENECONV, B – Bootscan, M – MaxChi, C – Chimaera, S – SiScan, T – 3Seq.

| Event | Recombinant Isolate | Genotype | Major Parent | Genotype | Minor Parent | Genotype | Detection Method |  |  |  |  |  |  |
| --- | --- | --- | --- | --- | --- | --- | --- | --- | --- | --- | --- | --- | --- |
|  |  |  |  |  |  |  | R | G | B | M | C | S | T |
| 1 | MH385014 | I | JN381831 | I | AY303798 | I | + | + |  | + | + | + | + |
| 2 | AF045551 | I | GU187972 | I | AY303798 | I | + | + | + | + | + | + | - |
| 3 | KU351668 | I | unknown |  |  |  | + | + | + | + | + | - | + |
| 4 | GU187972 | I | unknown |  |  |  | - | + | - | - | - | - | + |
| 5 | JF706282 | I | JN381851 | I | KT957423 | I | + | + | + | + | + | + | + |
| 6 | MH753128 | III | AF069076 | III | NC_001437 | I | + | + | + | + | + | - | + |
| 7 | KU351667 | I | KU508408 | I |  |  | + | + | + | + | + | + | + |
| 8 | KY885009 | III | AF069076 | III |  |  | + | + | + | + | + | + | + |
| 9 | JN381834 | I | JN381851 | I |  |  | + | + | - | + | + | + | + |
| 10 | AF098737 | III | JEVBEICG | III | unknown |  | + | + | + | - | - | - | + |
| 11 | JN381868 | III | JF706273 | III |  |  | + | + | - | + | + | + | + |
| 12 | KU871322 | III | KU871363 | III |  |  | + | + | + | - | - | - | + |
| 13 | JF706269 | III | JEVBEICG | III |  |  | + | + | + | - | - | - | + |
| 14 | JN381864 | III | JN381865 | III |  |  | - | - | - | + | + | + | + |
| 15 | KU508409 | I | MZ540901 | I |  |  | + | + | + | - | - | - | + |
| 16 | MZ702743 | III | KT229572 | I |  |  | - | - | - | + | + | - | - |
| 17 | JN381866 | III | JN381858 | III |  |  | + | + |  | + | + | - | + |
| 18 | AF098736 | III | unknown |  | JEVBEICG | III | + | - | - | + | - | + | - |

### Phylogenetic Relationships

Our phylogenetic analysis resolved the five previously identified genotypes of JEV with high confidence (Figure 1). As we did not identify any novel genotypes in our data, this analysis is largely confirmatory of prior studies (10, 26, 35) as we found the JEV genotypes fell into 5 reciprocally monophyletic clades in an ascending pattern of diversification from V – I (Figure 1). It is notable that Genotypes II, IV and V are represented by only one, seven and three isolates respectively despite the significant increase in size of the present dataset, indicating either a low prevalence of these genotypes in nature, or a considerable bias in the collection and public dissemination of JEV genomic data.

**Figure 1:**
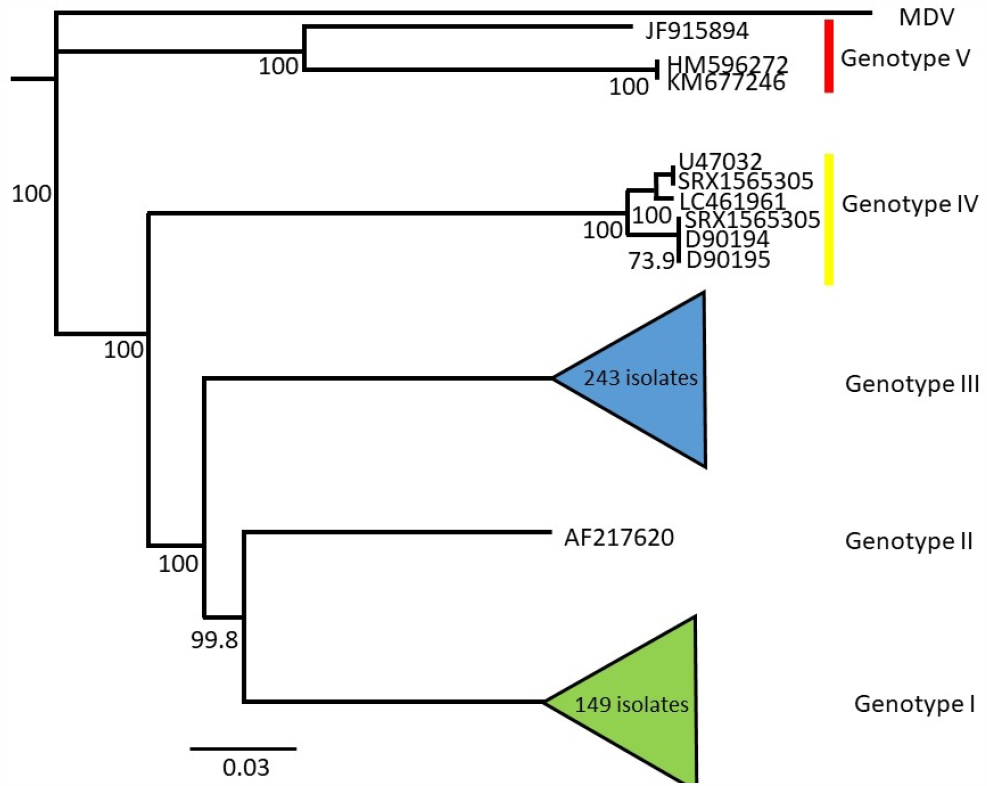
Phylogeny of JEV genome sequences, showing the five monophyletic genotypes of JEV. Branching order in the current analysis is confirmatory of prior studies of evolution of JEV. Node labels indicate bootstrap support, scale is in substitutions per site.

**Figure 2:**
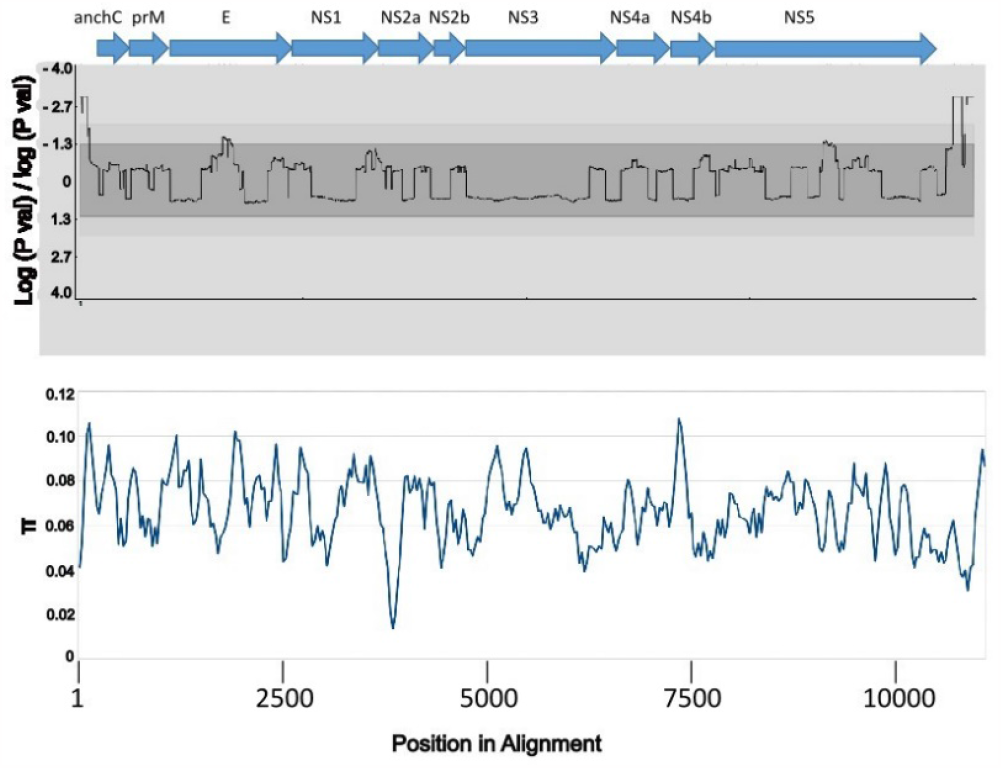
Recombination and selection analysis of JEV genomes. Top legend indicates the locations of protein coding genes, bottom legend indicates position in the genome alignment. Graph A depicts recombination break point probability per site, light grey indicates 99% probability, the mid grey area indicates 95% probability while the dark grey represents non-significant break points. Marginally significant break points were observed in structural protein E and Nonstructural protein 5. Graph B depicts nucleotide diversity (π) calculated using a sliding window analysis.

### Selection

Nucleotide diversity analysis showed relatively low rates of mutation across the genome (Fig 2, Graph B), with a notable decrease in π in NS2a and a notable increase in π in NS5. These rates of nucleotide diversity did not necessarily correspond with complimentary changes in Dn-Ds ratios (Table 2). All genes deviated significantly from a model of neutral evolution, with strongly negative Dn-Ds values indicative of a strong overall signal of purifying selection.

**Table 2:**
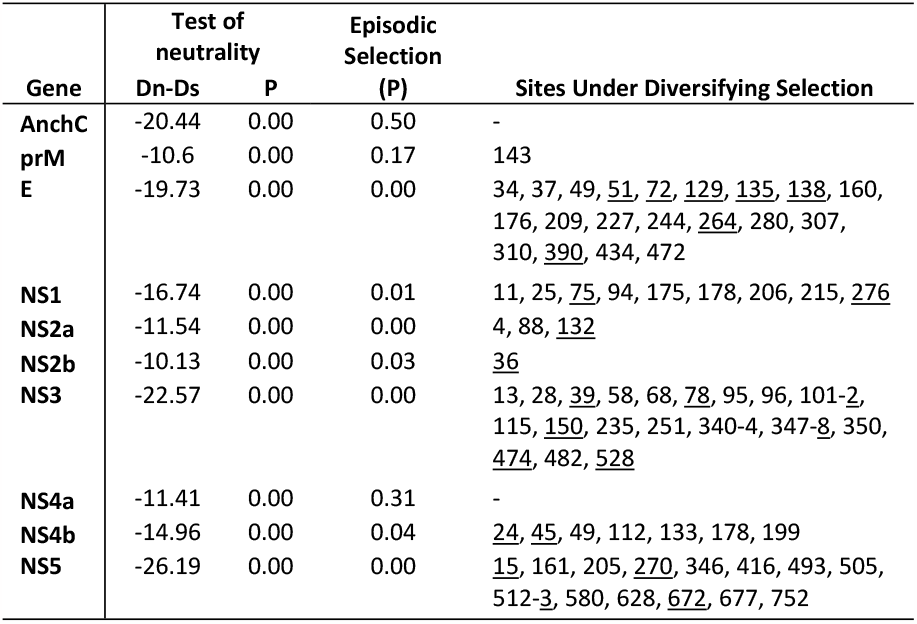
Results of selection analysis. Test of neutrality was conducted using an averaged codon based, two tailed Z test; Episodic selection was determined using a Branch-site Unrestricted Statistical Test for Episodic Selection; and sites under diversifying selection were determined using a Mixed Effects Model of Evolution analysis. Underlined sites are third position mutations.

| Gene | Test of neutrality |  | Episodic Selection (P) | Sites Under Diversifying Selection |
| --- | --- | --- | --- | --- |
|  | Dn-Ds | P |  |  |
| **AnchC** | -20.44 | 0.00 | 0.50 | - |
| **prM** | -10.6 | 0.00 | 0.17 | 143 |
| **E** | -19.73 | 0.00 | 0.00 | 34, 37, 49, <u>51</u>, <u>72</u>, <u>129</u>, <u>135</u>, <u>138</u>, 160, 176, 209, 227, 244, <u>264</u>, 280, 307, 310, <u>390</u>, 434, 472 |
| **NS1** | -16.74 | 0.00 | 0.01 | 11, 25, <u>75</u>, 94, 175, 178, 206, 215, <u>276</u> |
| **NS2a** | -11.54 | 0.00 | 0.00 | 4, 88, <u>132</u> |
| **NS2b** | -10.13 | 0.00 | 0.03 | <u>36</u> |
| **NS3** | -22.57 | 0.00 | 0.00 | 13, 28, <u>39</u>, 58, 68, <u>78</u>, 95, 96, 101-2, 115, <u>150</u>, 235, 251, 340-4, 347-8, 350, <u>474</u>, 482, <u>528</u> |
| **NS4a** | -11.41 | 0.00 | 0.31 | - |
| **NS4b** | -14.96 | 0.00 | 0.04 | <u>24</u>, <u>45</u>, 49, 112, 133, 178, 199 |
| **NS5** | -26.19 | 0.00 | 0.00 | <u>15</u>, 161, 205, <u>270</u>, 346, 416, 493, 505, 512-3, 580, 628, <u>672</u>, 677, 752 |

Branch-site Unrestricted Statistical Test for Episodic Selection (BUSTED)(49) analysis showed that some branches in the phylogenetic tree for the genes E, NS1, NS2a, NS2b, NS3, NS4b and NS5 all displayed episodic diversifying selection (Table 2). Within these genes, a Mixed Effects Model of Evolution (MEME) (50) showed that a small proportion of sites (0.7% of the genome) displayed episodic diversifying selection (Table 2) with the number of sites in each gene found to be under diversifying selection corresponding to the significance of the results of the BUSTED analysis. While summary statistic approaches demonstrate strong evolutionary conservation of genotypes at a genomic scale, more nuanced analysis shows that a small number of sites throughout the genome are diversifying in a manner likely to be adaptive. Of particular interest is the strong episodic diversifying selection of protein E – the major surface protein of the virion. Experimental evolutionary approaches have shown that specific mutations in surface proteins can generate thermo-tolerance in viruses (57, 58) and the mutations observed in the JEV E protein may assist in the evolution of this virus to persist in novel environments. Similarly, large numbers of sites under diversifying selection were observed in the large, multifunction, non-structural proteins NS3 and NS5, but due to their multi-functional nature, the specific selection pressures driving the diversification of these mutations will be difficult to determine.

## Conclusions

Our study conforms to the results of previous studies to demonstrate the five genotypes of JEV determined at the gene level are robust with increased sampling and whole genome phylogenetic analysis. We demonstrate that while recombination may occur at relatively small scales within JEV genotypes, it is unlikely to be a major driver of genomic diversity and evolution. We also show that despite widespread purifying selection acting on the JEV genome, there are a small number of sites under diversifying selection. Experimental approaches to determine the functional impact of these mutations are likely to yield insights into the evolutionary driving forces that precipitate geographic range, host and vector expansion in JEV.

## Acknowledgements

We thank Dr Richard Weir, Dr Rachel De Arajuo and Dr Vidya Bhardwaj for providing comments on the MS prior to submission. We thank the Australian Department of Agriculture, Fisheries and Forestry for funding (Grant C08586).

## Notes

### Competing Interest Statement

The authors have declared no competing interest.

